# PRODUCTION OF BIOETHANOL FROM WATER HYACINTH USING MONOCULTURE AND CO-CULTURE OF MICROORGANISMS

**DOI:** 10.1101/2024.09.03.611102

**Authors:** Idowu Samuel DADA, Titilayo FEMI-OLA, Opeyemi LASISI, Olalekan ADEOSUN

## Abstract

Water hyacinth is a lignocellulosic raw material for long-term suitable production of bioethanol. Though water hyacinth is considered to be a cause of ecological disorder, however, due to its inherent chemical composition consisting of higher cellulosic components, it may be proven to be a source for lignocellulosic ethanol and other value-added products. This study investigated bioethanol production from water hyacinth using fermentation using *Aspergillus niger*, *Saccharomyces cerevisiae*, and *Bacillus cereus*. The effect of the pH on the bioethanol fermentation was studied, also changes in reducing sugar content and ethanol concentration were determined. Fourier transform infrared spectrometry (FT-IR) was used to identify the functional groups of the bioactive component which measures the vibration of bonds in chemical functional groups for the samples obtained after distillation. The pH of the fermentation media for mono-culture and co-culture fermentation of water hyacinth with *Aspergillus niger*, *Saccharomyces cerevisiae*, and *Bacillus cereus* reduced from 6.0 to 3.7. There was an increase in the reduced sugar concentration during the fermentation period with the highest value (27%) obtained after 10 days of co-culture fermentation of water hyacinth with *A. niger* and *S. cerevisiae.* Co-culture fermented water hyacinth with *A. niger* and *S. cerevisiae* yielded the distillate with the highest ethanol concentration (35.4%). The FT-IR analysis of the distillate showed the presence of alcohol, aldehydes, and ketones.

## 1.0 Introduction

Water hyacinth (*Eichhornia crassipes*) is a water weed or aquatic plant that pollutes waterways and causes eutrophication, energy shortage, and endangers fish, among other issues (Jha and Namdeo, 2022). Solutions to address these problems include utilizing water hyacinth sustainably to produce renewable bioenergy, biofertilizer, and remedy contaminated water (Jha *et al*., 2024; Ilo *et al*., 2020). The potential for producing bioethanol through hydrolysis, fermentation, and distillation of water hyacinth has become popular over the years (Sharma *et al*., 2020). Factors that affect the production of bioethanol from water hyacinth include pretreatment, fermentation methods, and fermentation microbial strains (Deffar *et al*., 2024). This research is tested using monoculture and coculture of microbial strains *Aspergillus niger, Saccharomyces cerevisiae, and Bacillus cereus*. The development of new microbes that can efficiently break down cellulose and ferment sugars is crucial in the production of bioethanol with increased yield at a lower cost (Tse *et al*., 2021).

Energy is the driving force behind human development. As nonrenewable energy reserves are depleting, human civilization has entered an era of sustainable use of renewable energy (Mohammed *et al*., 2021). Bioethanol is one of the most important renewable energy sources and is produced from crops such as maize, corn, barley, sugar cane, and others (Jeevan Kumar *et al*., 2020). However, these crops have relatively low biomass content, making it economically unfeasible to use them for bioethanol production (Chen *et al*., 2021). The biotechnological method, incorporating the use of microbial strains, can produce bioethanol as certain microbes, such as yeast, can ferment sugars into bioethanol (Melendez *et al*., 2022). Therefore, lignocellulose is a potential raw material for bioethanol production. Lignocellulose is the most commonly used raw material in bioethanol production due to its advantages, such as large reserves and low cost (Pant and Kuila, 2022). Despite these advantages, the production of bioethanol from lignocellulose also faces several challenges that hinder its economic feasibility (Broda *et al*., 2022).

## 2.0 Materials and method

### 2.1 Sample Collection

Water hyacinth was used as raw material which was collected from a local pond at Ekiti State University, Ado-Ekiti, Ekiti State, Nigeria. The samples were brought to the Microbiology laboratory of Ekiti State University, Ado -Ekiti for studies on bioethanol production.

### 2.2 Physical Pretreatment

In the laboratory, the water hyacinth was thoroughly washed several times with tap water to remove the adhering dirt. The roots were discarded as they have been reported to absorb heavy metal pollutants from water bodies and dried under the sun for about 48 hours. It was then chopped into small pieces of size 1 - 2 cm and dried again at 70 °C in a hot air oven for 1 h. It is further grinded to even smaller particles with mortar and pestle of size 2 mm. The grinded water hyacinth was stored in a dry container at room temperature and subjected to acid pretreatment (Tirva *et al*., 2022).

### 2.3 Acid Pretreatment

Acid pretreatment was carried out in conical flasks (250ml) each by mixing 50g of dried water hyacinth (Biomass) with 20ml of 2% (v/v) sulfuric acid (H_2_SO_4_). After that, the flasks were autoclaved at 121°C for 15 minutes and further cooled down to room temperature. The pretreated samples were dried in an oven at 65°C (Gyalai-Korpos *et al.,* 2011). Pretreated samples were subjected to fermentation (Tirva *et al*., 2022)

### 2.4 Microorganism and its Maintenance

Pure culture of *Bacillus cereus, Aspergillus niger, and Saccharomyces cerevisiae* was obtained from the stock culture in the Microbiology Laboratory at Ekiti State University, Ado Ekiti. *Bacillus cereus* was maintained on Nutrient agar slants at 4°C. *Aspergillus niger* and *Saccharomyces cerevisiae* were maintained on Potatoes Dextrose Agar (PDA) slants at 4°C.

### 2.5 Submerged Fermentation

Fermentation was carried out in pre-sterilized 1000ml capacity bottles containing 50g each of pretreated powered water hyacinth in 500 ml of distilled water. The pH of the medium was adjusted to 6.0 using 1N HCL and 1N NaOH. The flasks were sterilized by autoclaving at 121 °C for 15 minutes. The water hyacinth first flask for coculture fermentation was inoculated with 5 ml *Aspergillus niger* broth and *Saccharomyces cerevisiae* broth (CAS) while the second flask for coculture fermentation was inoculated with 5 ml *Bacillus cereus* broth and *Saccharomyces cerevisiae* broth (CBS). For mono-culture fermentation, monoculture fermentation was sequentially inoculated. The first flask for mono-culture fermentation was inoculated, with 5ml *Aspergillus niger* broth (MA), while the second flask for monoculture fermentation was inoculated with 5 ml *Bacillus cereus* broth (MB). The flasks were incubated at ambient temperatures on an orbital shaker set at 250 rpm for 10 days (Zhang *et al*., 2016)

### 2.6 Determination of Reducing Sugar During Fermentation

Spectrophotometry was applied to find the concentration of reducing sugar in each sample. From the sample, 5ml was been withdrawn. Into 5ml of each sample, 2ml of Benedict’s reagent was added. The resulting mixture was placed in a boiling water bath for 5 minutes. Positive results gave rise to brown coloration. The Absorption was read at 447 nm. These readings were used to plot a graph of the concentration against days. The blank which is 10 ml distilled water and 2 ml Benedict’s reagent was used to zero the spectrophotometer (Tirva *et al*., 2022).

### 2.7 Determination of Ethanol Concentration Produced During Fermentation

Ethanol concentrations from the fermentation were determined daily for 10 days. 1 ml of ethanol sample was poured into a test tube, 7 ml of distilled water was added, and 2 ml of acidified potassium dichromate was added. It was then heated for 40 minutes in a water bath. The absorbance was read at 580nm using a UV-visible spectrophotometer as described by Mojovic*et al*. (2009).

### 2.8 Centrifugation and Distillation

After fermentation, the broth was centrifuged at 6000rpm for 10 minutes. The supernatant was collected and fed into a simple laboratory distillation column. The boiling temperature of ethanol is 78 °C hence distillation was carried out around that temperature to facilitate the evaporation of ethanol. The vapor was collected and condensed using cold water to circulate the column. The distillate with ethanol was recovered in a flask at the other end of the column.

### 2.9 Fourier Transform Infrared Spectrometry (FT-IR) Analysis

Fourier Transform Infra-Red Spectroscopy (FT-IR), was done for the samples obtained after distillation in which maximum bioethanol production was expected to check if peaks representing ethanol bonding were present or not. Infra-red (IR) analysis was done with the aid of an infra-red spectrophotometer (Perkin-Elmer, spectrumbx) at the Federal University of Oye-Ekiti, Ekiti State Nigeria. A drop of distillate was placed on a fused sodium chloride (NaCl) cell. It was carefully placed on a cell clamped loosely and fixed on the infra-red beam. The infra-red data was compared to the table of IR frequencies using the methods of Johnson *et al*. (2012).

## 3.0 RESULTS

### 3.1 Effect of pH on Bioethanol Production

Fermentation was carried out with an initial pH of 6. The changes in pH during co-culture and monoculture fermentation of water hyacinth are shown in Figures 1-2. Figure 1 showed the pH result of the co-culture of water hyacinth during fermentation with a broth of both *Aspergillus niger* and *Saccharomyces cerevisiae* (CAS), the pH remains constant on day 2 (6.0) followed by a progressive decrease in pH value on day 4 (5.8) day 6 (5.7) day 8 (5.0) day 10 (4.8). Co-culture of water hyacinth during fermentation of water hyacinth by both *Bacillus cereus* and *Saccharomyces cerevisiae* (CBS), the pH value decreased from day 2 (5.9) followed by a progressive decrease on day 4 (5.7), day 6 (5.6) day 8 (5.4) till day 10 (5.0).

**Figure 1:**
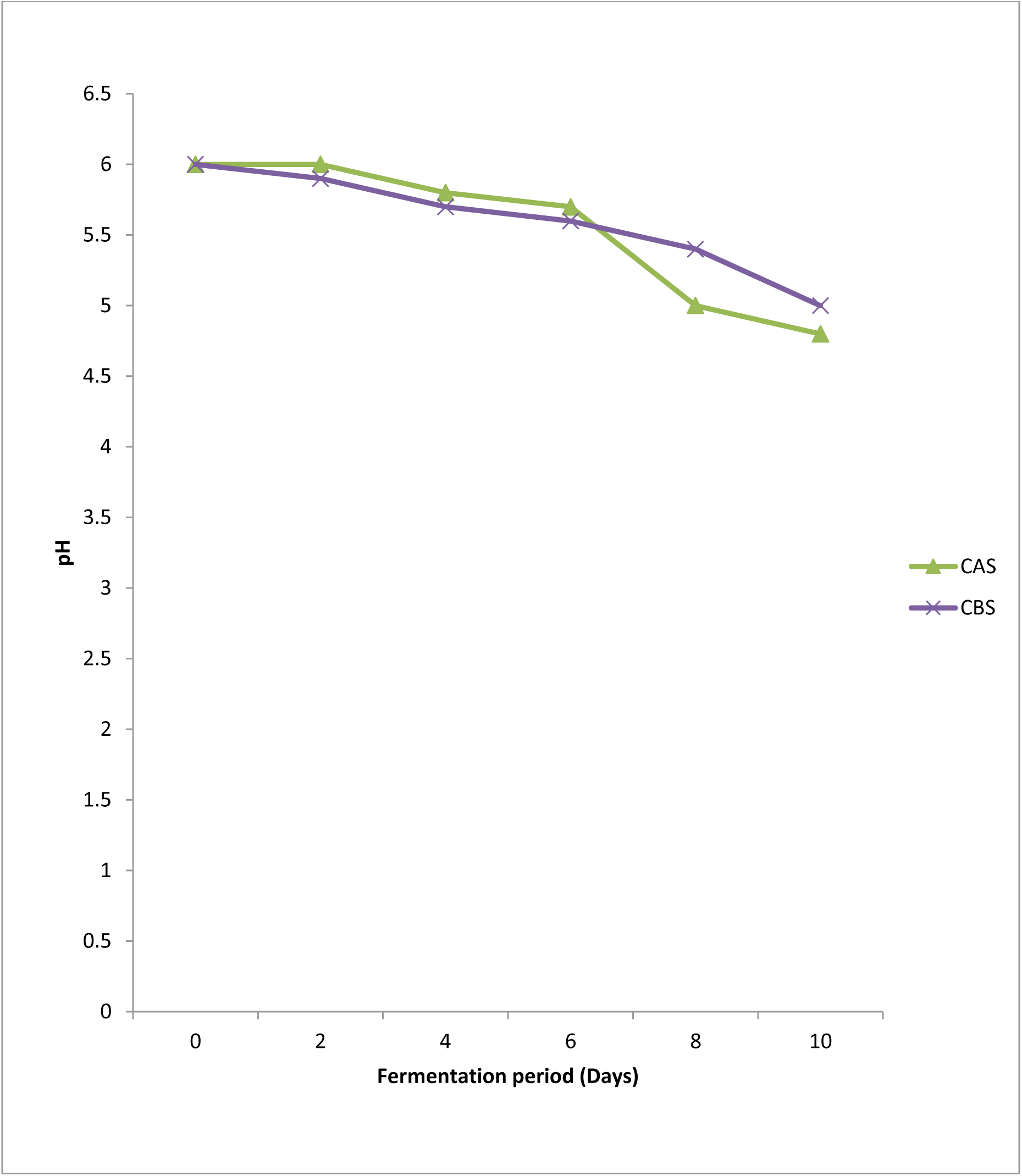
pH of coculture of water hyacinth during fermentation of the samples. Key: Water hyacinth (CAS: *Aspergillus niger* and *Saccharomyces cerevisiae*, CBS: *Bacillus cereus* and *Saccharomyces cerevisiae*).

**Figure 2:**
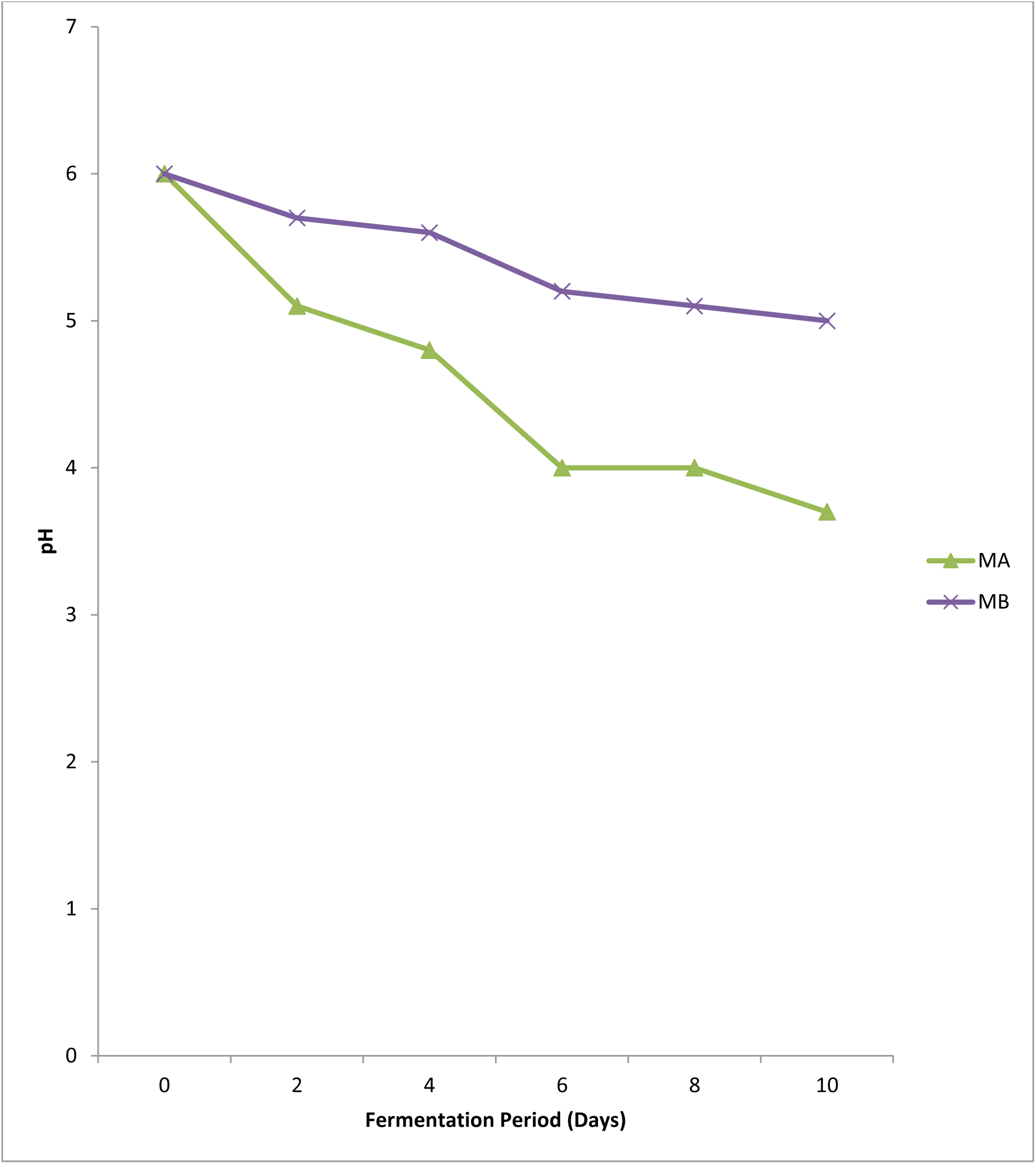
pH of monoculture of water hyacinth during fermentation of the samples. Key: Water hyacinth (MA: *Aspergillus niger*, MB: *Bacillus cereus*).

The changes in pH during mono-culture fermentation of the water hyacinth by *Aspergillus niger* (MA) and *Bacillus cereus* (MB) are shown in Figure 2. For monoculture fermentation with *Aspergillus niger* (MA), a progressive decrease was observed in pH value from day 2 (5.1), day 4 (4.8), day 6 (4.0), day 8 (4.0) till day 10 (3.7). The pH of mono-culture during fermentation of water hyacinth with *Bacillus cereus* (MB), the pH value decreased on day 2 (5.7), followed by generally decreased in day 4 (5.6), day 6 (5.2), day 8 (5.3), till day 10 (5.0).

### 3.2 Changes in Reducing Sugar Content during Fermentation

Figure 4 shows the result of reducing sugar of coculture of water hyacinth during fermentation of the samples. Coculture of water hyacinth during fermentation of both *Aspergillus niger* and *Saccharomyces cerevisiae* (CAS) on day 8 showed the highest concentration of reducing sugar (27%) and coculture of water hyacinth during fermentation of both *Bacillus cereus* and *Saccharomyces cerevisiae* (CBS) in day 8 showed the highest concentration of reducing sugar (20%).

**Figure 3:**
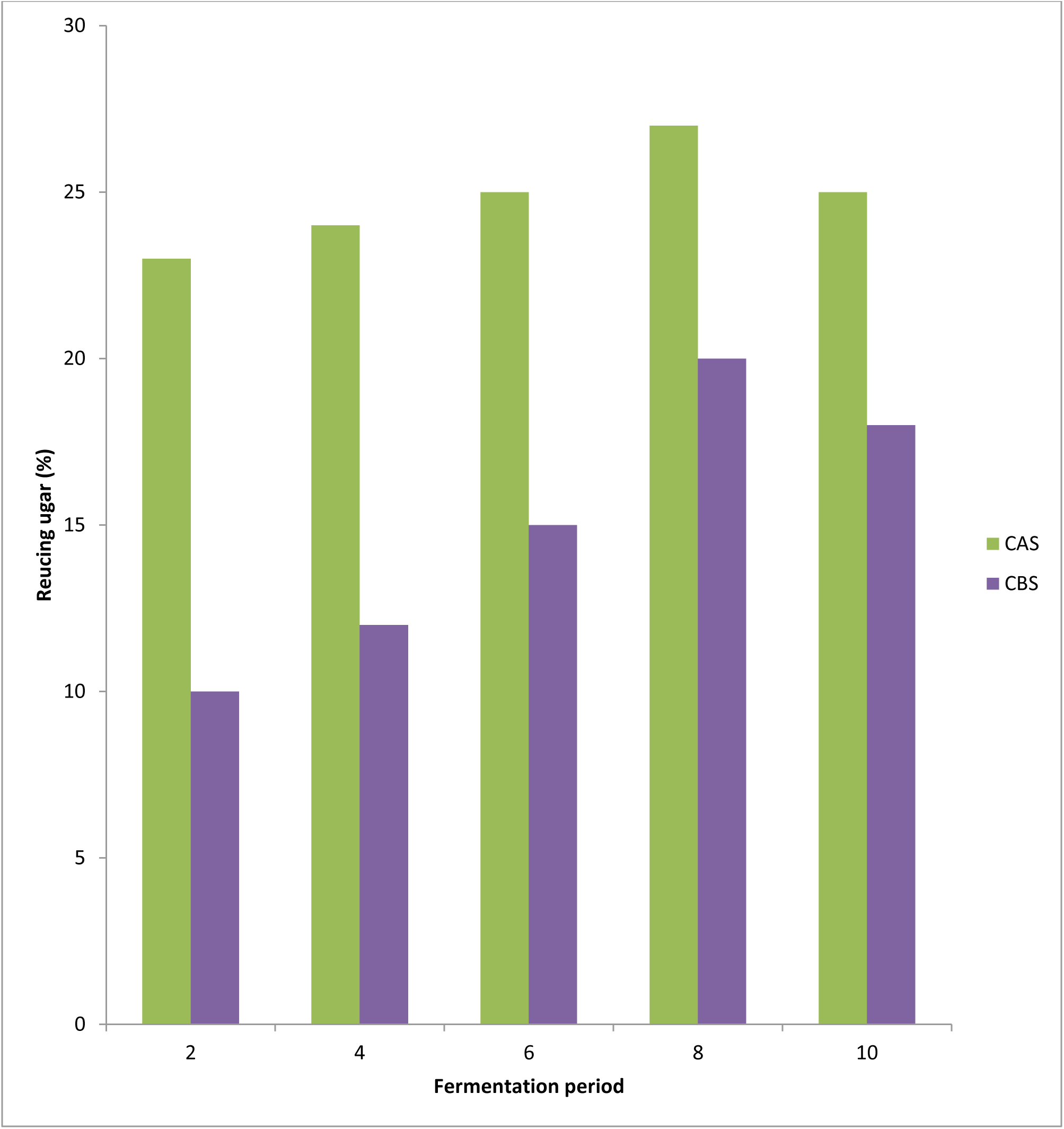
Reducing sugar of coculture water hyacinth during fermentation of the samples. Key: Water hyacinth (CAS: *Aspergillus niger* and *Saccharomyces cerevisiae*, CBS: *Bacillus cereus* and *Saccharomyces cerevisiae*).

**Figure 4:**
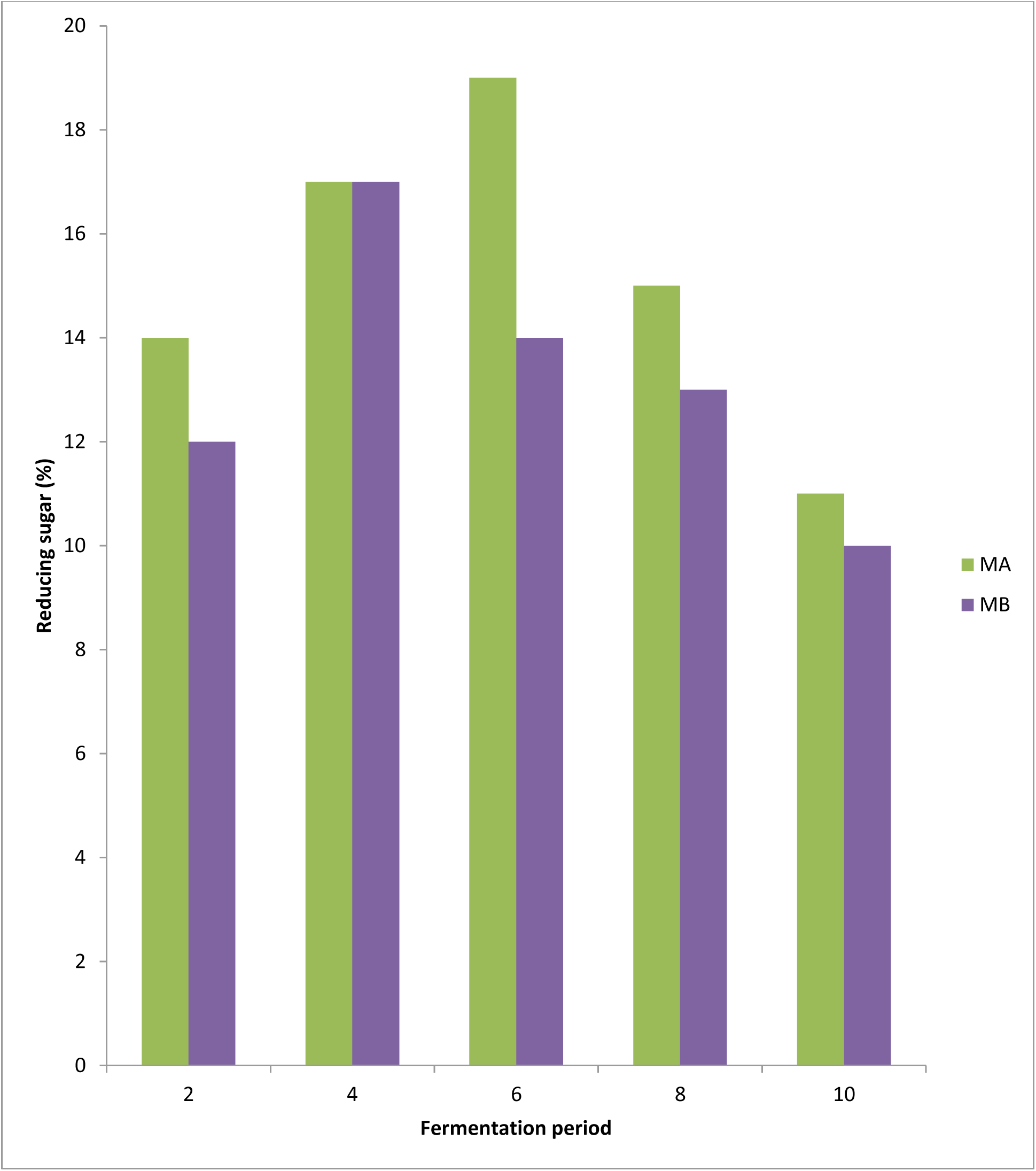
Reducing sugar of monoculture of water hyacinth during fermentation of the samples. Key: Water hyacinth (MA: *Aspergillus niger*, MB: *Bacillus* cereus).

Figure 5 shows reducing sugar of monoculture of water hyacinth during fermentation. Monoculture of water hyacinth during fermentation of *Aspergillus niger* (MA) on day 3 showed the highest concentration of reducing sugar (19%) while monoculture of water hyacinth during fermentation of *Bacillus cereus* (MB) on day 8 showed the highest concentration of reducing sugar (17%).

**Figure 5:**
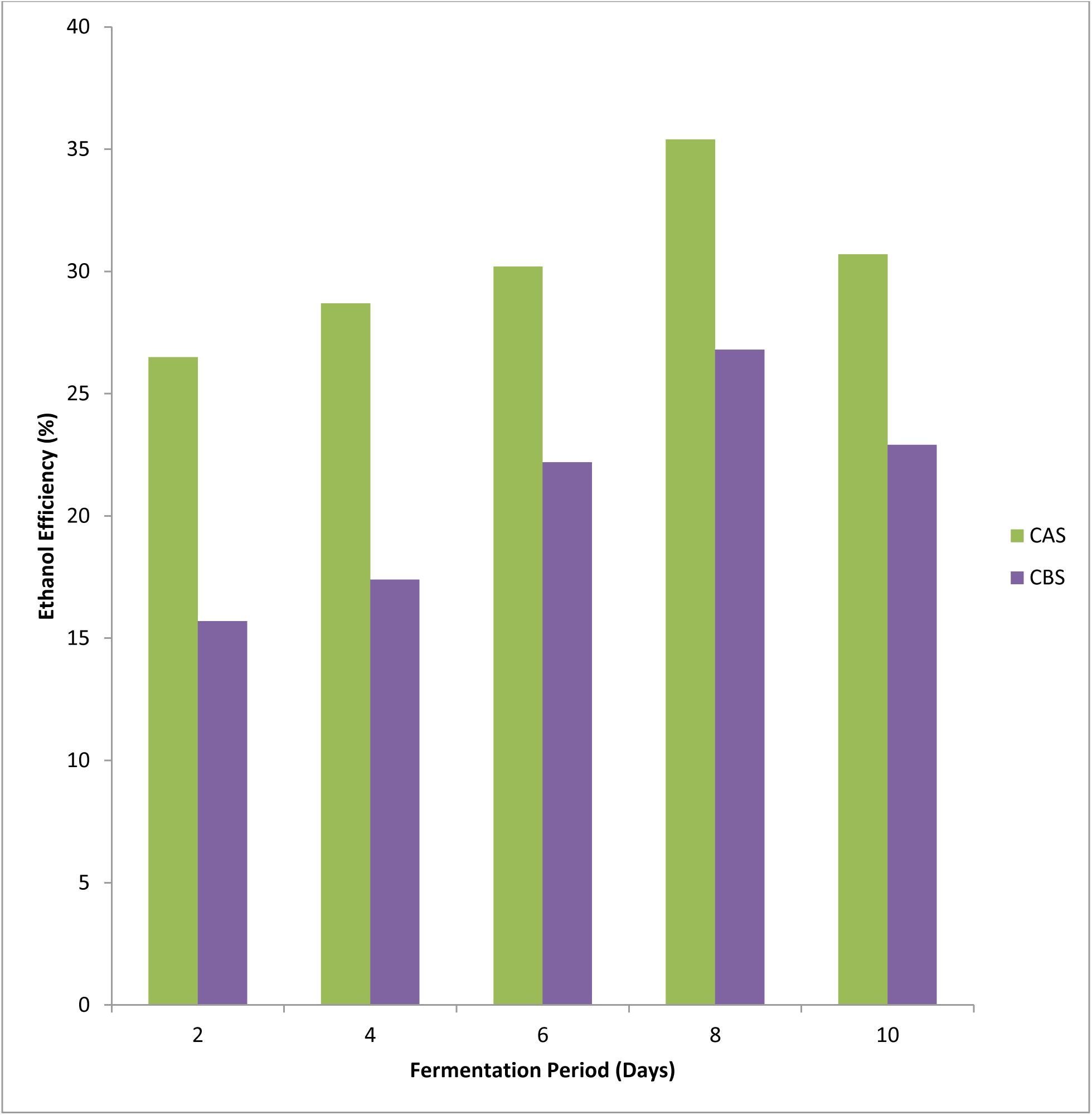
Ethanol concentration of coculture of water hyacinth during fermentation of the samples. Key: Water hyacinth (CAS: *Aspergillus niger* and *Saccharomyces cerevisiae*, CBS: *Bacillus cereus* and *Saccharomyces cerevisiae*).

**Figure 6:**
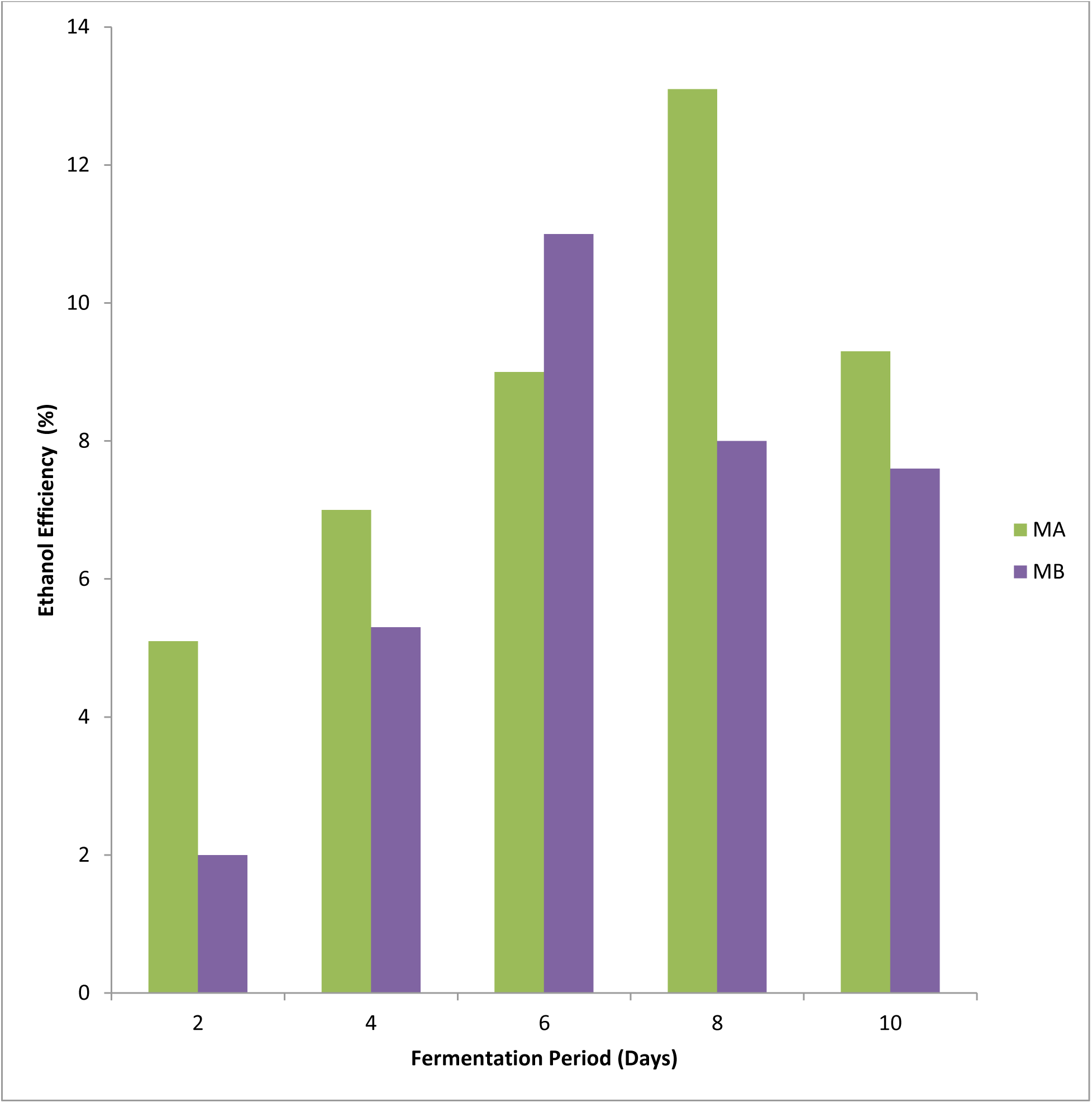
Ethanol concentration of monoculture of water hyacinth during fermentation of the samples. Key: Water weed (MA: *Aspergillus niger*, MB: *Bacillus cereus*).

### 3.3 Determination of Ethanol Concentration

The results of ethanol yield shown from coculture of water hyacinth with *Aspergillus niger* and *Saccharomyces cerevisiae* (CAS) and coculture of water hyacinth with *Bacillus cereus* and *Saccharomyces cerevisiae* (CBS) showed a maximum ethanol yield in day 6 fermentation period in each of the substrates. The peak ethanol production using *Aspergillus niger* and *Saccharomyces cerevisiae* (CAS) was 35.4% followed by 30%, 30.7%, 28.7%, and 26.5% and *Bacillus cereus* and *Saccharomyces cerevisiae* (CBS) had 26.8% followed by 22.9%, 22.2%, 17.4%, and 15.7%.

For monoculture of water hyacinth with *Aspergillus niger* (MA) showed a maximum ethanol yield on day 6 and the monoculture of water hyacinth of the *Bacillus cereus* (MB) showed a maximum ethanol yield in the day 4 fermentation period of the substrates. The peak ethanol production using *Aspergillus niger* (MA) was 13.1% followed by 9.3%, 9%, 7%, and 5.1% while *Bacillus cereus* (MB) had 11% followed by 7.6%, 8%, 5.3%, and 2%.

### 3.4 Fourier-Transform Infrared Spectroscopy (FT-IR) Analyses

The FT-IR analysis of the distillates of coculture of water hyacinth after fermentation of the samples using *Aspergillus niger* and *Saccharomyces cerevisiae* (CAS).

Figure 8, revealed three characteristic peaks. The observed absorptions ranged from strong and weak absorption bands. The absorption bands at 3300.48 cm^-1^ was assigned to the presence of O-H stretching vibration. The bands at 1634.45 cm^-1^ are assigned to C–O. Table 2 shows the distillate of coculture of water hyacinth after fermentation of the samples using *Bacillus cereus* and *Saccharomyces cerevisiae* (CBS). Figure 8 revealed four characteristic peaks. The observed absorptions ranged from strong and weak absorption bands. The absorption bands at 3333.03 cm^-1^ were assigned to the presence of O-H stretching vibration. The peak at 2202.33 and 2020.24 cm^-1^ was assigned to C-N. The presence of ketones and Aldehydes was confirmed by the absorption of 1634.12 cm^-1^.

Distillate of monoculture of water hyacinth after fermentation of the samples using *Aspergillus niger* (MA). Figure 12, revealed three characteristic peaks which are interpreted in Table 7. The observed absorptions ranged from strong and weak absorption bands. The absorption bands at 3333.64 cm^-1^ were assigned to the presence of O-H stretching vibration. The presence of ketones and Aldehydes was confirmed by the absorption 1634.04 cm^-1^. The distillate of monoculture of water hyacinth after fermentation of the samples with *Bacillus cereus* (MB) in Figure 13, revealed four characteristic peaks which are interpreted in Table 8. The observed absorptions ranged from strong, and weak absorption bands. The absorption bands at 3330.00 cm^-1^ were assigned to the presence of O-H stretching vibration. The peak at 2212.61 cm^-1^ was assigned to C-N. The presence of ketones and Aldehydes was confirmed by the absorption 1633.75 cm^-1^.

**Figure 7:**
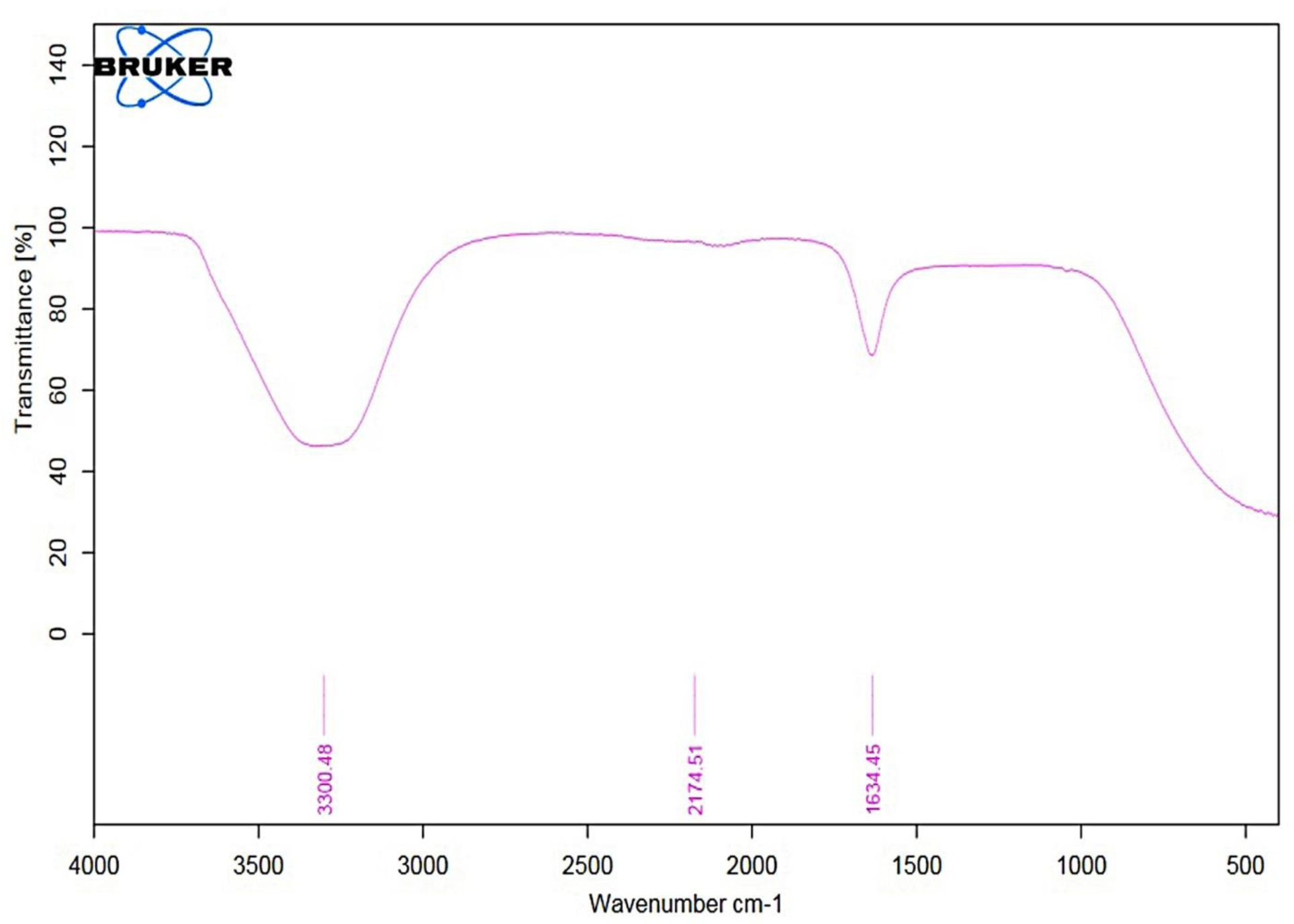
FTIR Spectrum of distillate of coculture of water hyacinth after fermentation of the samples using *Aspergillus niger* and *Saccharomyces cerevisiae* (CAS).

**Table 1:**
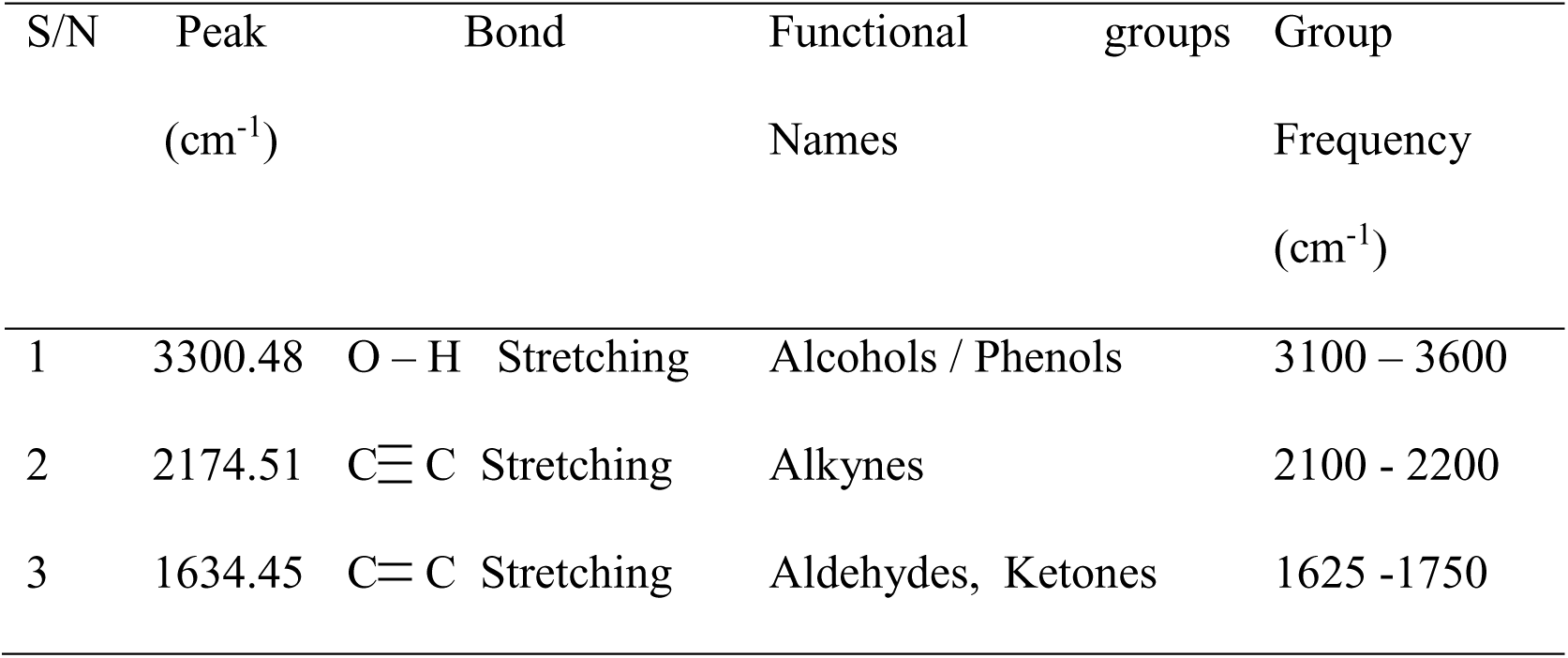
Distillate of coculture of water hyacinth after fermentation of the samples using *Aspergillus niger* and *Saccharomyces cerevisiae* (CAS)

**Figure 8:**
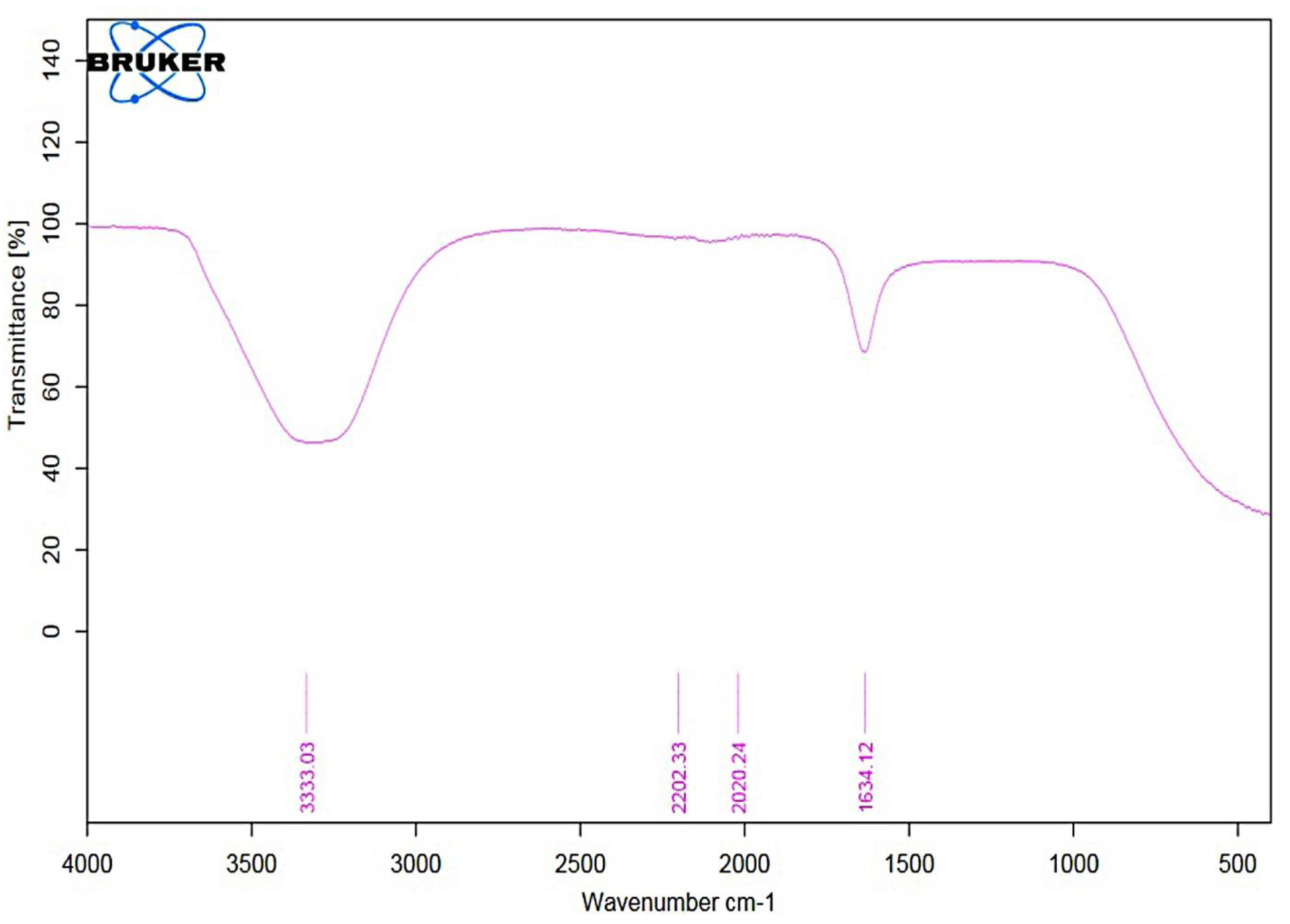
FTIR Spectrum of distillate of coculture of water hyacinth after fermentation of the samples using *Bacillus cereus* and *Saccharomyces cerevisiae* (CBS).

**Table 2:**
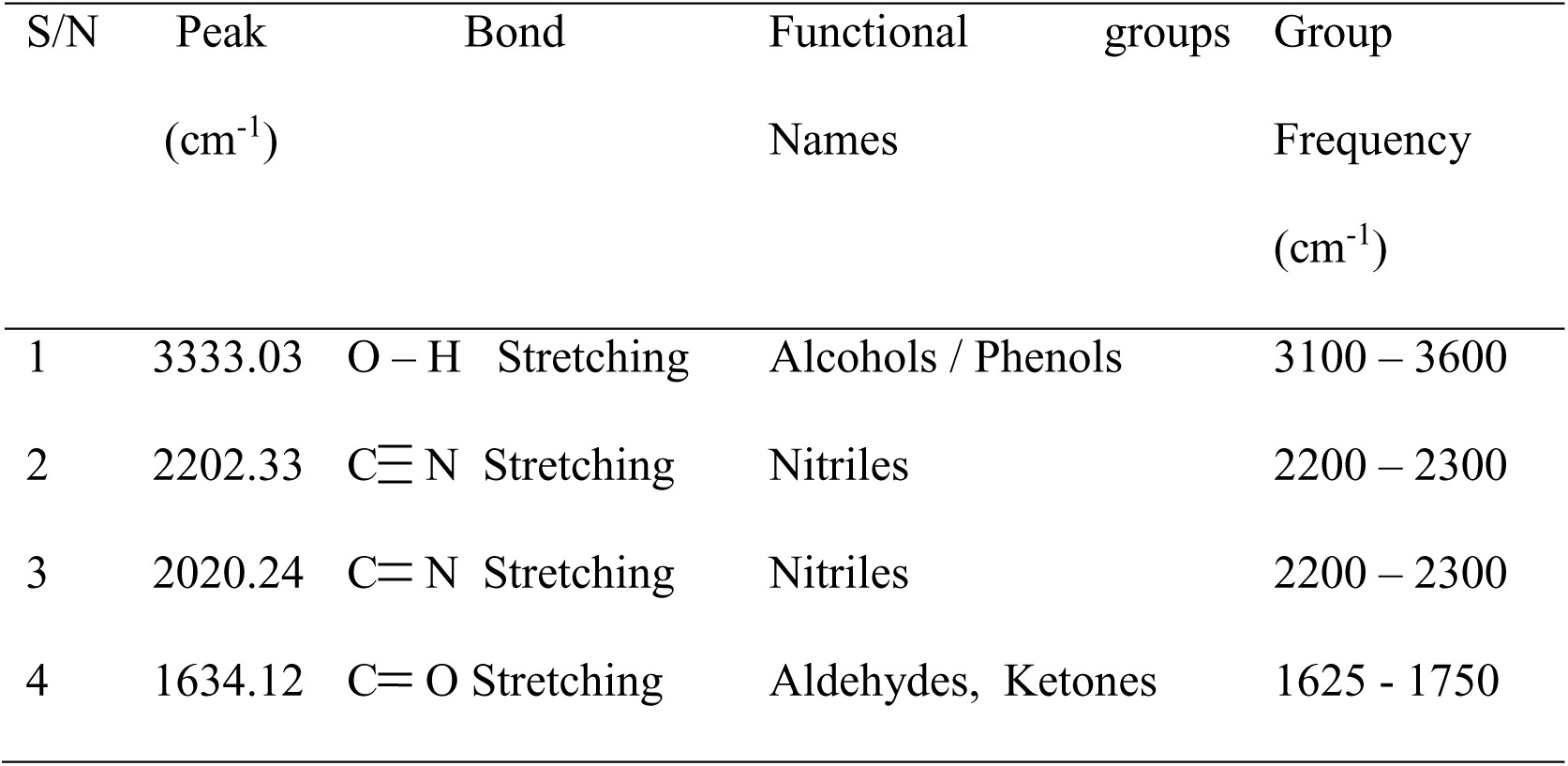
Distillate of coculture of water hyacinth after fermentation of the samples using *Bacillus cereus* and *Saccharomyces cerevisiae* (CBS)

**Figure 9:**
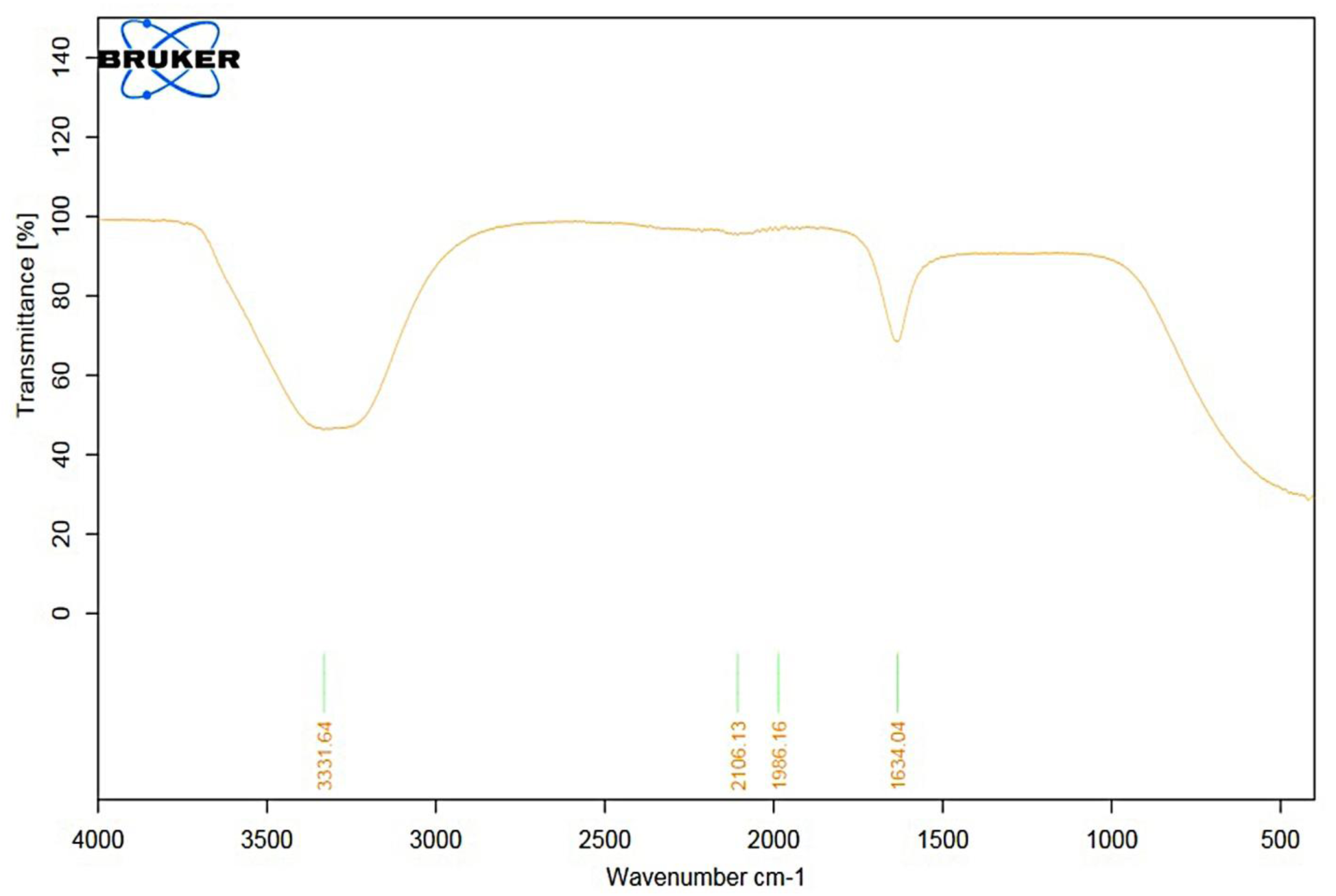
FTIR Spectrum of distillate of monoculture of water hyacinth after fermentation of the samples *Aspergillus niger* (MA).

**Table 3:**
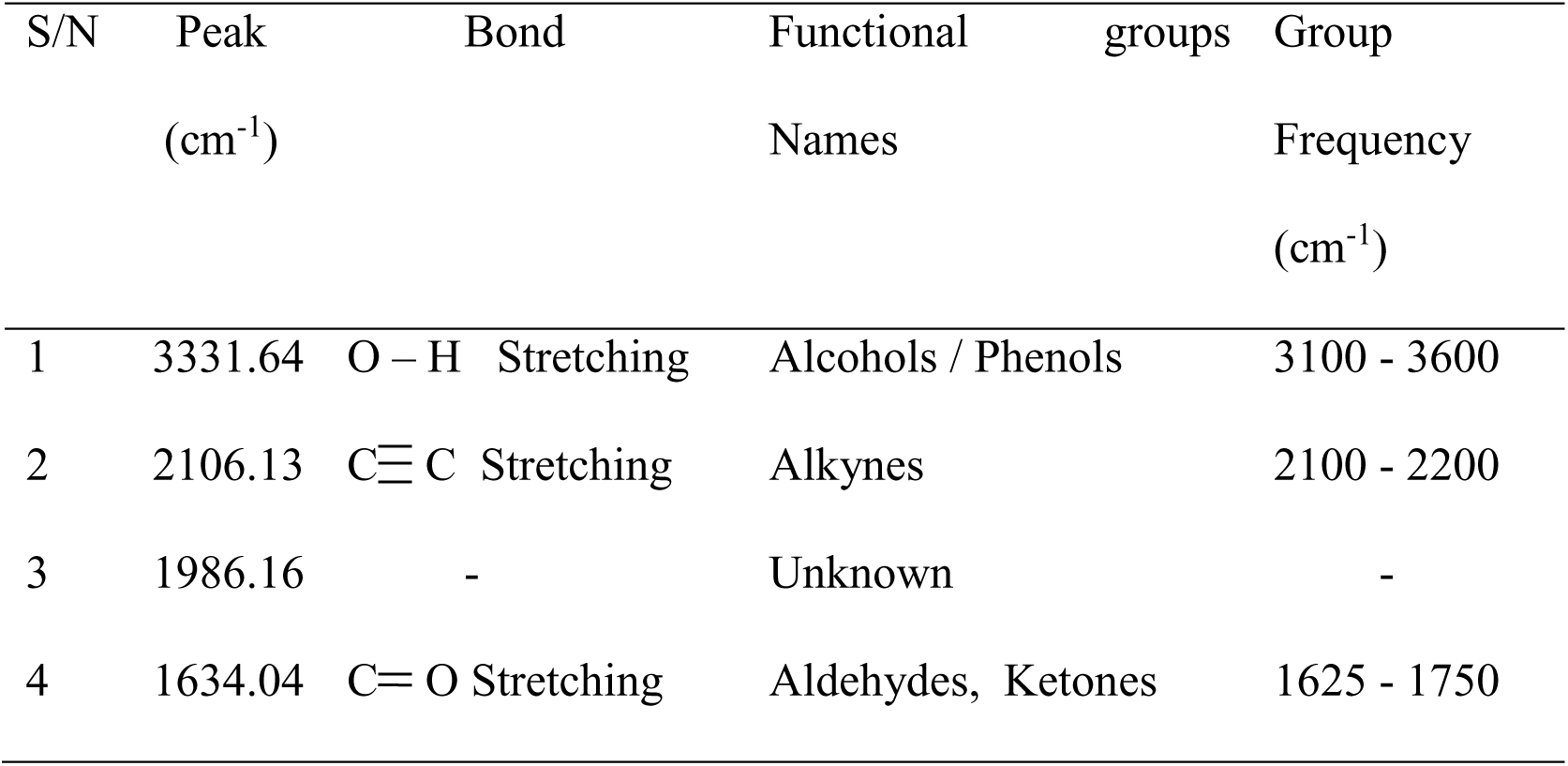
Distillate of monoculture of water hyacinth after fermentation of the samples *Aspergillus niger* (MA).

**Figure 10:**
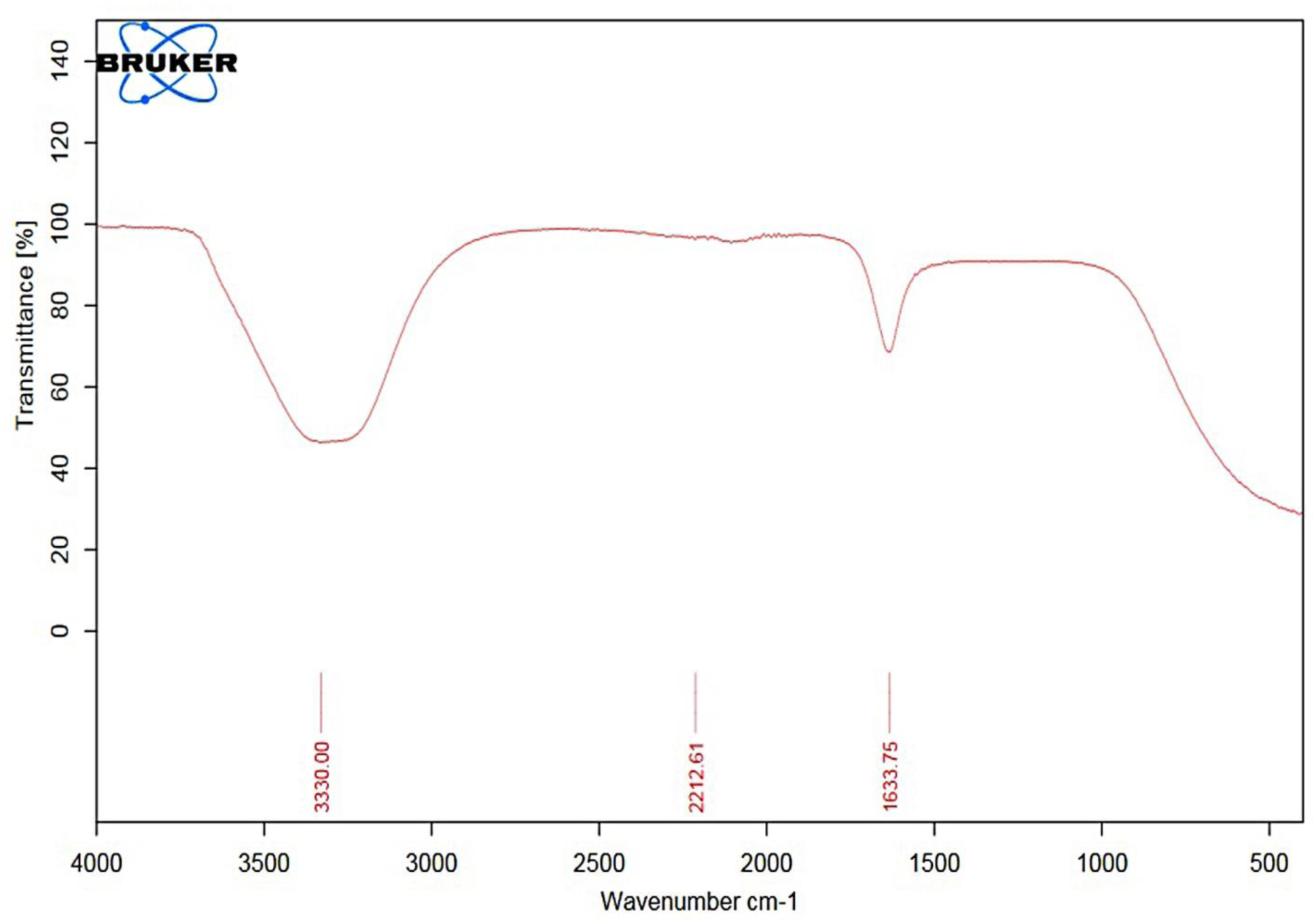
FTIR Spectrum of monoculture of water hyacinth after fermentation of the samples *Bacillus cereus* (MB).

**Table 4:**
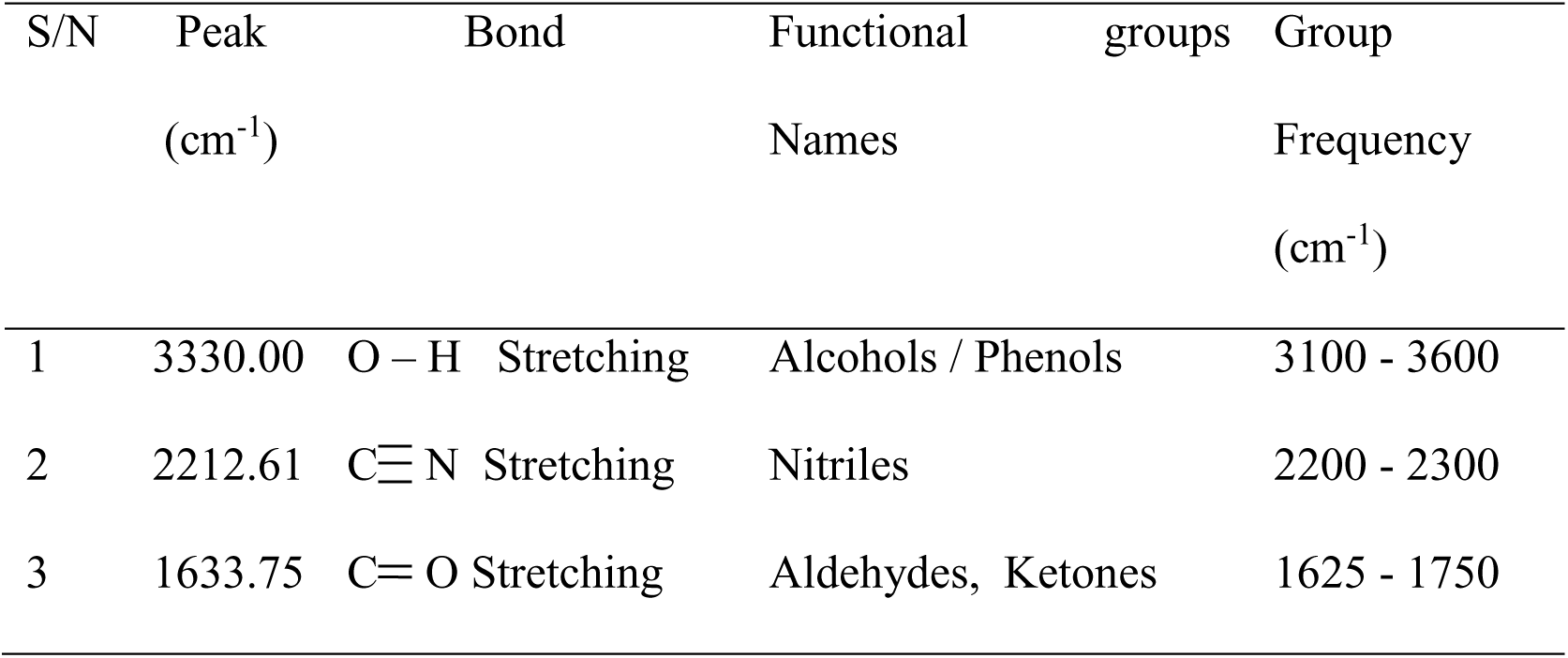
Monoculture of water hyacinth after fermentation of the samples *Bacillus cereus* (MB).

## 4.0 DISCUSSION

This study evaluated the efficiency of *Aspergillus niger* and *Saccharomyces cerevisiae*, *Bacillus cereus,* and *Saccharomyces cerevisiae* for coculture fermentation and monoculture of *Aspergillus niger* only and *Bacillus cereus* only for water hyacinth. This study reports that the pH of the broth of *Aspergillus niger* and *Saccharomyces cerevisiae* (CAS) and *Bacillus cereus* and *Saccharomyces cerevisiae* (CBS) for coculture fermentation of water hyacinth generally decreased during the days of the fermentation period. This co-relates with the work of Braide *et al*. (2016).

The decrease in pH value maybe due to an indication that the fermentation process became more acidic as a result of the production of other secondary metabolites and activities of microorganisms in the fermentation medium. For monoculture of *Aspergillus niger* (MA) and *Bacillus cereus* (MB) of water hyacinth. There was also a decrease in pH value. As the pH decreases, the fermenting broth becomes more acidic, thus changing the metabolic activities of the microorganisms for increased ethanol production.

Reducing sugar of coculture of water hyacinth during fermentation of both *Aspergillus niger* and *Saccharomyces cerevisiae* (CAS) in day 8 showed the highest concentration of reducing sugar (27%) than reducing sugar of both *Bacillus cereus* and *Saccharomyces cerevisiae* (CBS) in day 8 (20%). Where similar observations were made by Ganguly *et al*. (2013).

The decrease in sugar level on the fourth day could be attributed to the decrease in the sugar present in the broth was fermented to alcohol and other by-products such as glycerol and CO_2_ (Bekatorou, *et al*., 2006). Reducing sugar of water hyacinth for monoculture of *Aspergillus niger* (MA) in day 6 has the highest concentration of reducing sugar (19%) than *Bacillus cereus* (MB) in day 4 had (17%). The decrease in sugar level on the second day could be attributed to the decrease in the sugar present in the broth was fermented to alcohol. Bacteria are known to multiply faster than yeast. *Bacillus cereus* might reach the lag phase earlier than *Aspergillus niger* and therefore utilized its substrate faster (Randive *et al.,* 2015).

The result shows the highest ethanol concentration obtained on the 8^th^ day for both coculture of water hyacinth of *Aspergillus niger* and *Saccharomyces cerevisiae* (CAS), *Bacillus cereus* and *Saccharomyces cerevisiae* (CBS). This could be attributed to the fact that yeast cells are probably at their lag phase trying to synthesized necessary enzymes for their metabolism thereby converting the readily available sugar to ethanol. The highest concentration may represent the maximum tolerable limit of ethanol by the yeast cells. Ethanol has been reported as a well-known inhibitor of the growth of microorganism due to its effect on the mitochondrial of yeast cell and some enzyme such as hexokinase and dehydrogenase. Never the less, some strains of the yeast *Saccharomyces cerevisiae* how tolerance and can adapt to high concentration of ethanol (Wang *et al.,* 2014).

Monoculture of water hyacinth of *Aspergillus niger* (MA) reaching the peak in day 8 of fermentation and then declined (Wang *et al.,* 2014) and broth of the *Bacillus cereus* (MB) showed a maximum ethanol yield in day 6 fermentation period of the substrates.

This could be attributed to the fact that bacterium was progressing to the stationary phase and could no longer utilize the limited sugar present in the samples (Wang *et al.,* 2014). The absorption bands at 3333.03 cm^-1^ was assigned to the presence of O-H stretching vibration, the absorption in this region was due to the alcoholic or phenolic compounds present in the co-culture of water hyacinth after fermentation of the samples using *Aspergillus niger* and *Saccharomyces cerevisiae*, (CAS) which confirmed the presence ethanol by the absorption 3333.03 cm^-1^ (Ashokkumar and Ramaswamy, 2014). The band at 1634.68 cm^-1^ are assigned to C-O which possibly due to the presence of ketones, and Aldehydes. The absorption bands at 3300.84 cm^-1^ was assigned to the presence of O-H stretching vibration. The absorption in this region was due to the alcoholic or phenolic compounds present in the coculture of water hyacinth after fermentation of the samples using *Bacillus cereus* and *Saccharomyces cerevisiae*, (CBS) which confirmed the presence ethanol by the absorption 3333.03 cm^-1^ (Ashokkumar and Ramaswamy, 2014). The peak at 2202.33 cm^-1^ and 2020.24 cm^-1^ was assigned to C-N. The presence of ketones, and Aldehydes was confirmed by the absorption 1634.12 cm^-1^.

The absorption bands at 3333.64 cm^-1^ was assigned to the presence of O-H stretching vibration, the absorption in this region was due to the alcoholic or phenolic compounds present in the monoculture of water hyacinth after fermentation of the samples by *Aspergillus niger* and *Saccharomyces cerevisiae* (MA), which confirmed the presence ethanol by the absorption 3286.80 cm^-1^ (Ashokkumar and Ramaswamy, 2014). The presence of ketones, and Aldehydes was confirmed by the absorption 1634.04 cm^-1^. The absorption bands at 3332.38 cm^-1^ was assigned to the presence of O-H stretching vibration, the absorption in this region was due to the alcoholic or phenolic compounds present in the monoculture of water hyacinth after fermentation of the samples *Bacillus cereus* and *Saccharomyces cerevisiae* (MB) which confirmed the presence ethanol by the absorption 3330.00 cm^-1^ (Ashokkumar and Ramaswamy, 2014). The peak at 2212.61 cm^-1^ was assigned to C-N which indicates the presence of Nitriles compound. The presence of ketones, and Aldehydes was confirmed by the absorption 1633.75 cm^-1^.

## CONCLUSION

It can be concluded that waterweed materials can be fermented for the production of bioethanol. Water hyacinths are rich in cellulose and thus can be considered as a suitable substrate for the production of bioethanol. This study also revealed that *Aspergillus niger*, *Saccharomyces cerevisiae,* and *Bacillus cereus* serve as a better microorganism for bioethanol production in which a high yield of bioethanol was produced in this study. Water hyacinth was a suitable substrate for bioethanol production by *Aspergillus niger* and *Saccharomyces cerevisiae*, high yield of ethanol was produced. This substrate could therefore be used for large-scale bioethanol production. Therefore, the findings of this work suggest that ethanol can be produced from water hyacinth rather than allowing it to contribute a nuisance to the environment.

## Declarations

### Ethics approval and consent to participate

Not applicable

### Consent to publication

Not applicable

### Availability of Data and Materials

The data sets used and/or analyzed during this study are available from the corresponding author upon reasonable request.

### Code Availability

Not applicable

### Authors’ contribution

DADA, Idowu Samuel *, FEMI-OLA, Titilayo, LASISI, Opeyemi, ADEOSUN, Olalekan. FEMI-OLA, Titilayo, conceived the project and defined research methodology; DADA, Idowu Samuel, LASISI, Opeyemi, ADEOSUN, Olalekan performed the experiments. DADA, Idowu Samuel, LASISI, Opeyemi, ADEOSUN, Olalekan wrote the paper.

## Acknowledgement

We thank the Department of Microbiology and Chemistry Research Laboratory for their technical support.

